# Methane-linked mechanisms of electron uptake from cathodes by *Methanosarcina barkeri*

**DOI:** 10.1101/415653

**Authors:** Annette R. Rowe, Shuai Xu, Emily Gardel, Arpita Bose, Peter Girguis, Jan P. Amend, Mohamed Y. El-Naggar

## Abstract

The *Methanosarcinales*, a lineage of cytochrome-containing methanogens, have recently been proposed to participate in direct extracellular electron transfer interactions within syntrophic communities. To shed light on this phenomenon, we applied electrochemical techniques to measure electron uptake from cathodes by *Methanosarcina barkeri*, which is an important model organism that is genetically tractable and utilizes a wide range of substrates for methanogenesis. Here we confirm the ability of *M. barkeri* to perform electron uptake from cathodes and show that this cathodic current is linked to quantitative increases in methane production. The underlying mechanisms we identified include, but are not limited to, a recently proposed association between cathodes and methanogen-derived extracellular enzymes (e.g. hydrogenases) that can facilitate current generation through the formation of reduced and diffusible methanogenic substrates (e.g. hydrogen). However, after minimizing the contributions of such extracellular enzymes and using a mutant lacking hydrogenases, we observe a lower potential hydrogen-independent pathway that facilitates cathodic activity coupled to methane production in *M. barkeri*. Our electrochemical measurements of wild-type and mutant strains point to a novel and extracellular-enzyme-free mode of electron uptake able to take up electrons at potentials lower than - 498 mV vs. SHE (over 100 mV more reduced than the observed hydrogenase midpoint potential under these conditions). These results suggest that *M. barkeri* can perform multiple modes (hydrogenase-mediated and free extracellular enzyme-independent) of electrode interactions on cathodes including a mechanism pointing to a direct interaction, which has significant applied and ecological implications.

**Importance:** Methanogenic Archaea are of fundamental applied and environmental relevance. This is largely due to their activities in a wide range of anaerobic environments, generating gaseous reduced carbon that can be utilized as a fuel source. While the bioenergetics of a wide variety of methanogens has been well studied with respect to soluble substrates, mechanistic understanding of their interaction with solid phase redox active compounds is limited. This work provides insight into solid phase redox interactions in *Methanosarcina* using electrochemical methods. We highlight a previously undescribed mode of electron uptake from cathodes, that is potentially informative of direct interspecies electron transfer interactions in the *Methanosarcinales*.

## Introduction

The ability of microbes to generate methane, a major component of natural gas and biogas from anaerobic digestion, as well as a powerful greenhouse gas, has important technological and environmental implications. The formation of methane through the reduction of carbon dioxide (CO_2_) is mediated by a distinct group of Euryarchaeota (1). The energy captured during this process reflects one of life’s lowest energy yielding metabolisms (2). Hydrogen (H_2_), the predominant electron donor for methanogenesis, is strikingly close in redox potential to CO_2_ (the primary terminal electron acceptor) under standard conditions, which illustrates the relatively slim energetic yields often available to methanogens (3). As such, these organisms employ a variety of ecologic and biochemical strategies – some newly and/or poorly understood – in order to persist in the wide range of anoxic environments, including some of the deepest and hottest ecosystems on Earth (4, 5).

One or more steps in the methanogenesis pathway coupling CO_2_ reduction from H_2_ (and in some cases formate) oxidation can be endergonic (energy consuming) under physiologic conditions (6). The initial reduction step, from CO_2_ to a formyl-methanofuran intermediate (6), requires ferredoxin—an iron-sulfur electron carrier with a lower redox potential than H_2_ in most biological contexts (1). Two strategies are known for (re)generating reduced ferredoxin in methanogens (7). The first strategy, utilized by the lineage that does not contain cytochromes, couples the first ferredoxin requiring step (energetically unfavorable) with the final reduction of a methyl group bound to Coenzyme M to produce methane and regenerate the Coenzyme M – Coenzyme B intermediate (energetically favorable) (8, 9). This occurs via a flavin-mediated electron-bifurcating enzyme complex (8, 9). The second strategy, utilized by methanogenic clades that contain cytochromes, employs an energy converting hydrogenase (Ech) that utilizes a cellular ion motive force to drive the energetically up-hill reduction of ferredoxin from H_2_(7). The biochemical differences represented in these two strategies underlie the ecologic trade-offs between respiration rate (allowing faster substrate utilization) and energetic yield (maximizing efficiency of energy capture), which often dictate the differential success of these groups under various competitive environmental conditions.

In many environmental systems, hydrogenotrophic methanogens also face energetic challenges with respect to electron donor limitation, given that environmental H_2_ concentrations are commonly near or below the energetic threshold for methanogenesis (3, 6). It has been suggested that H_2_ is often of syntrophic origin in methanogenic environments. Syntrophic interactions involve a metabolic intermediate (i.e. hydrogen) generated via fermentation at low hydrogen partial pressures by one organism being consumed by a partnering microbe (i.e. methanogen) (10). The energetic favorability of syntrophic fermentations is stabilized by the subsequent consumption of fermentation intermediates by the syntrophic partners, thereby maintaining favorable environmental conditions. The first such example was the co-culture of an ethanol fermenting bacterium and a hydrogenotrophic methanogen (11). Though this metabolic strategy is thought to dominate in many anoxic environments, the prevalence of hydrogen and/or formate as a metabolic intermediate has been widely debated (12, 13). It has proved challenging to match the known rates of diffusion for these intermediates with the necessary concentrations that would reflect energetic favorability for both the fermenter and methanogen syntrophic partners. Several studies have suggested that the rates of diffusion are too slow at syntrophically relevant concentrations to account for the rates of methanogenesis that are observed in many environmental systems (14). Put simply, a faster means of electron transfer is likely necessary to explain the measured rates.

A previously demonstrated co-culture of the ethanol-fermenting *Geobacter metallireducens* and methane-producing *Methanosarcina barkeri* highlighted the potential for a hydrogen-free and formate-free mode of interspecies electron transfer (15). The ability of *G. metallireducens* to perform direct extracellular electron transfer (EET) to solid surfaces is well known (16). Since mutants impaired in this direct EET were incapable of forming viable ethanol-consuming methane-generating consortia (15), it was suggested that these co-cultures share electrons through direct interspecies electron transfer (DIET) (17). However, while the mechanistic basis of direct outward EET from *Geobacter* species is well studied, there is currently no proposed mechanism for such a direct inward EET mechanism into the methanogenic partners.

To address this knowledge gap, we investigated the potential of *M. barkeri* to interact with solid phase electron sources (i.e. cathodes) as a surrogate for obtaining electrons from a syntrophic partner. To date, the only known mode of cathodic electron uptake by methanogens is catalyzed by cell-derived free enzymes (predominantly hydrogenases) that can attach to electrodes (18). In that case, methane is generated from electrochemically produced electron donors, such as H_2_ or formate (18). Notably, these findings were obtained with *Methanococcus maripaludis*—an organism in the clade of methanogens without cytochromes. We hypothesized that a previously unknown mode of electron transfer may be present in cytochrome containing methanogens, and that this mode may be distinguished electrochemically. Here, we apply electrochemical techniques to investigate the potential for direct electron uptake by *M. barkeri*—a cytochrome containing methanogen previously shown to generate methane in consortia with *G. metallireducens*. This has potentially important implication for understanding both the ecologic trade-offs between direct and hydrogen-based syntrophic partnerships as well as the potential utility of these organisms in bio-electrode technologies.

## Results

### M. barkeri generates methane in electrochemical cells with poised potentials lower than -400 mV SHE

We tested both methane production and electron uptake (cathodic current) by *M. barkeri* cell cultures in 3-electrode H-cell electrochemical reactors, with the working carbon cloth electrode poised between -400 mV and -500 mV vs. SHE, under two different culture conditions. The first consisted of cells in their growth or spent medium (growth cultures), and the second represented washed cells where the pre-grown cultures were centrifuged and resuspended in fresh basal medium lacking electron donor, reductants, vitamins or minerals (washed cultures). The second condition was chosen to mitigate the potential effects of the growth or spent media containing free enzymes (e.g. hydrogenases) capable of attaching to carbon electrodes and potentially masking cell-electrode interactions with enzyme-electrode reactions (as described in (18)). Compared to open circuit controls, *M. barkeri* cell cultures demonstrated increased methane production and yielded cathodic currents (Table 1). Both current and methane production were larger in preparations that included spent media (i.e. growth cultures), but both poised potential electrode experiments (i.e. growth and washed cultures) produced more methane than the open circuit controls (Table 1, Figure 1). Notably, the coulombic efficiencies (percentage of electrons that could be accounted for in the methane produced) were significantly larger in washed culture incubations compared to the growth cultures (Table 1). As previously suggested (18), this may be due to the influence of free hydrogenases in the spent media interacting with electrodes, resulting in a hydrogen pool that accounts for a portion of the coulombs drawn from electrode, but not converted to methane. To test this hypothesis, we performed cyclic voltammetry (CV) to determine patterns of electron uptake relative to the redox potentials in the aforementioned experiments.

**Table 1.**
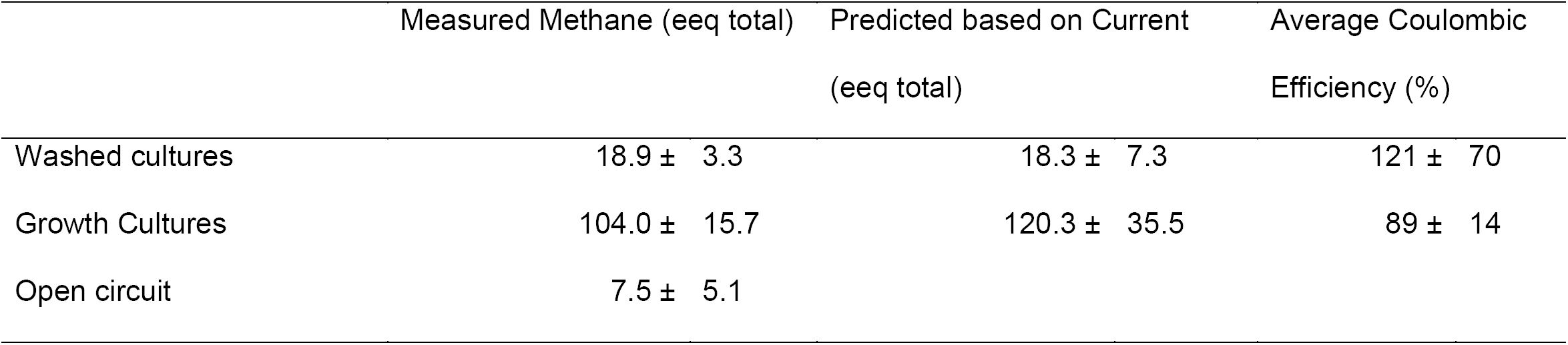
Methane generated and current consumption measured in wild-type *M. barkeri* experiments expressed in terms of electron equivalents (eeq) or moles of electron per experiment depicted in Figure 1. Each value represents at least experimental triplicates and error represents standard deviations.

**Figure 1.**
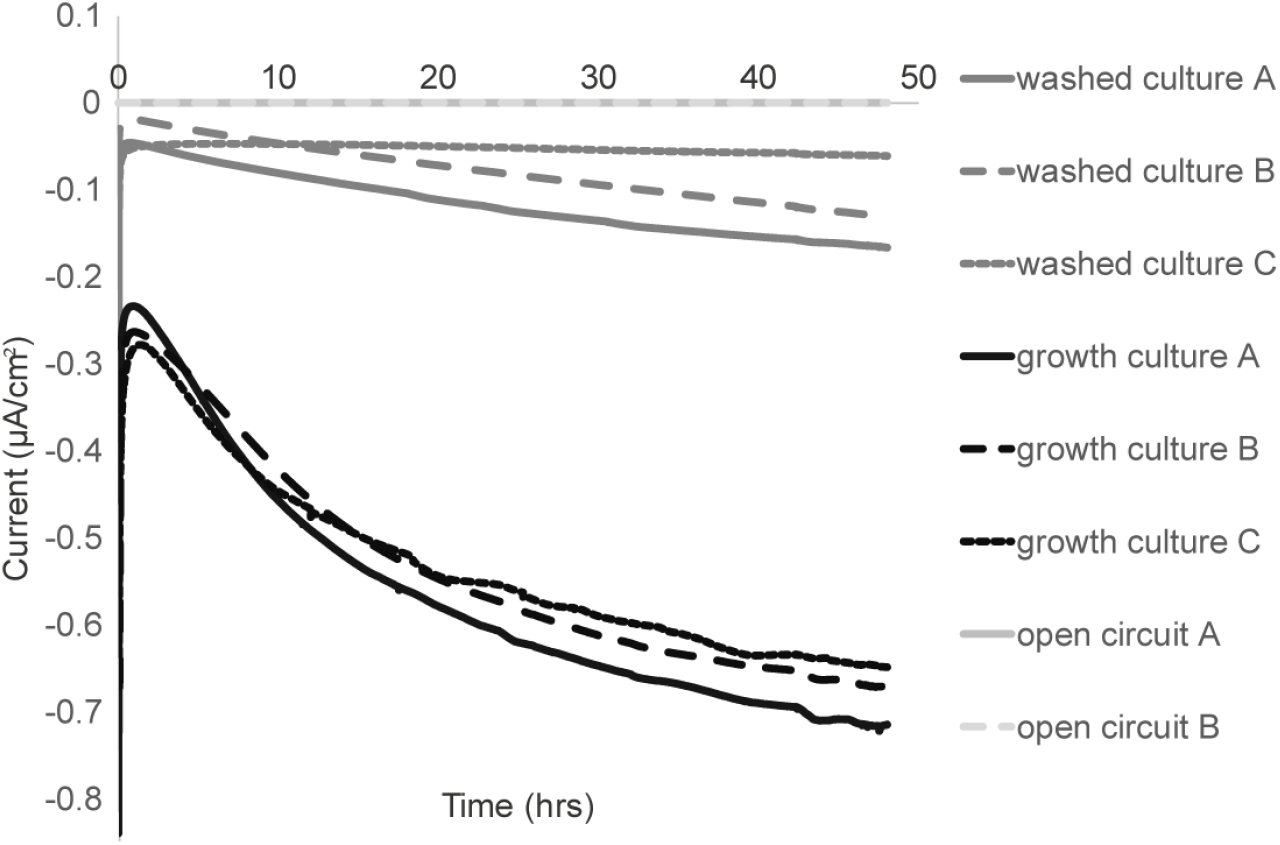
Electron uptake occurs in *M. barkeri* (MB) with and without growth media present at -450 mV. Comparison of current consumption in *M. barkeri* cells on poised electrodes. This includes experiments where: 1) the growth media was removed (washed culture); 2) cells were added along with the former growth media (growth culture); or 3) cells were washed to remove growth media but no potential was poised on the electrode (open circuit).

### Multiple electron uptake features observed with cyclic voltammetry

Cyclic voltammograms demonstrated two distinct catalytic features observed over the described experimental conditions. One catalytic feature was observed in both growth culture and washed culture experiments. A second, more positive catalytic wave was observed in only the growth culture incubations (which included potential free enzymes) (Figure 2A). The shared and predominant cathodic feature, as demonstrated by the first derivatives of the cathodic sweep of cyclic voltammograms, peaked at -498 mV to -509 mV SHE for growth cultures and washed cultures, respectively (Figure 2B). In the growth culture experiments, the first derivative analysis revealed other subtle features at higher redox potentials (-207 ± 5.6 mV and -303.7 ± 2.3 mV, Figure 2B). The -303 mV feature is consistent with the second catalytic feature observed and shares the redox potential for H_2_ production at pH 6.5-7 in this high salt system (range 380-300 mV depending on ionic strength). Notably, the corresponding feature observed in the first derivative of the anodic sweep (-296.7 ± 5.1 mV, Supplemental Figure S1A), and the small offset between the anodic and cathodic peaks is consistent with the activity of an electrode-attached enzyme (19, 20). This is in contrast to the ∼100 mV separation measured for the shared electron up-take feature (-509 ± 9.4 mV and -604 ± 13.3 mV for the anodic and cathodic sweeps) (Supplemental Figure S1B), which is consistent with previous observations of the more distant cell-electrode interactions (21–23). The - 207 mV diffusion-limited electrochemical feature observed is consistent with the redox indicator added to the growth media – resazurin –which has been observed electrochemically in growth media without cells, but not the basal media used in electrochemical experiments.

**Figure 2.**
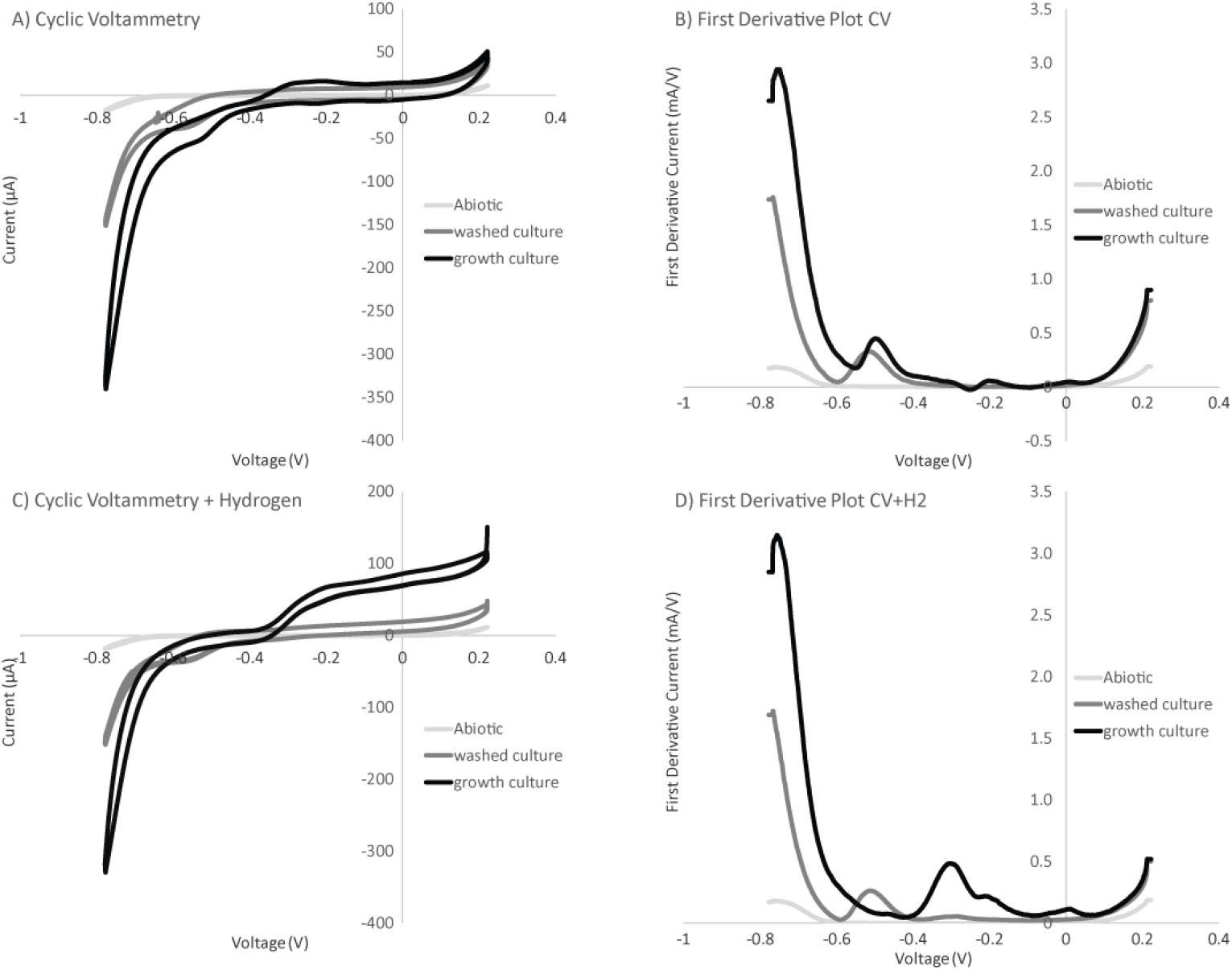
Differences observed in electron uptake features between washed cultures and growth culture experiments. Cyclic voltammetry (1 mV/sec scan rate over a -800 to 200 mV range) comparing voltage and current relationships between poised potential (-450 mV) experiments (shown in Figure 1) (A). First derivative plot of the cathode sweep (positive voltage to negative voltage) for each CV comparing peaks of electron uptake features observed (B). Replacing the headspace with a hydrogen and carbon dioxide gas mix (80% and 20% respectively) demonstrates some features are sensitive to hydrogen concentration (C). First derivative plot of the cathode sweep (positive voltage to negative voltage) for each hydrogen addition CV demonstrates the changes in features sensitive to hydrogen (D).

To determine if one or both CV features are indicative of hydrogenase activity, the same electrochemical reactors were purged with an 80/20 H_2_/CO_2_ gas mix to assess the impact of increasing H_2_ concentration. Given that hydrogenases are reversible enzymes, we expected that hydrogen would provide a substrate for oxidation by the enzymes, generating anodic current and inhibiting electron uptake. As predicted, a dramatic anodic catalytic wave was observed at the predicted hydrogenase redox potential in growth cultures when H_2_concentrations were greater than 1 mM (Figure 2C). This wave is consistent with hydrogenase-driven H_2_ oxidation at redox potentials above -405 mV SHE (onset potential observed in the anodic sweep of CV from Figure 2C). Neither catalytic activity (Figure 2C) nor a shift in redox potential of the predominant -509 mV feature (Fig. 2D) was significantly altered by increasing H_2_ in the washed cell experiments. Conversely, hydrogen greatly affected the putative hydrogenase feature in the growth culture experiments, which dramatically increased and dominated both the CV and first derivative plots of the cathode sweep in these experiments (Figure 2C-D). Taken collectively, the impact of different cell preparations and H_2_ addition suggest the presence of multiple electron uptake mechanisms in these *M. barkeri* electrochemical systems, including both hydrogen-dependent (hydrogenase-mediated) and hydrogen-independent (non-hydrogenase mediated) mechanisms.

### Current generation observed in M. barkeri hydrogenase deletion mutant with electrodes poised below -400 mV

To confirm the impact of a non-hydrogenase mediated mechanism of electron uptake that can be utilized for methane formation, electrochemical experiments were performed using a hydrogenase deletion mutant of *M. barkeri* (24). This mutant lacks the genes for all three types of hydrogenase present in *M. barkeri*: the ferredoxin-dependent energy converting hydrogenase (Ech), the cytoplasmic F_420_-dependent hydrogenase (Frh), and the periplasmic methanophenazine-dependent hydrogenase (Vht). Furthermore, this mutant does not grow with hydrogen as an electron donor or via acetate disproportionation. However, this strain is capable of growth via methanol disproportionation (24). Cathodic currents (i.e. electron uptake) were generated with this mutant in three-electrode systems with the working electrodes poised from -400 mV to - 500 mV (Figure 3A-B). The coulombic efficiencies of current linked to methane production for washed preparations of the hydrogenase mutant were comparable (180 ± 90 %) to those observed in the wild-type washed culture preparations (Table 1).

**Figure 3.**
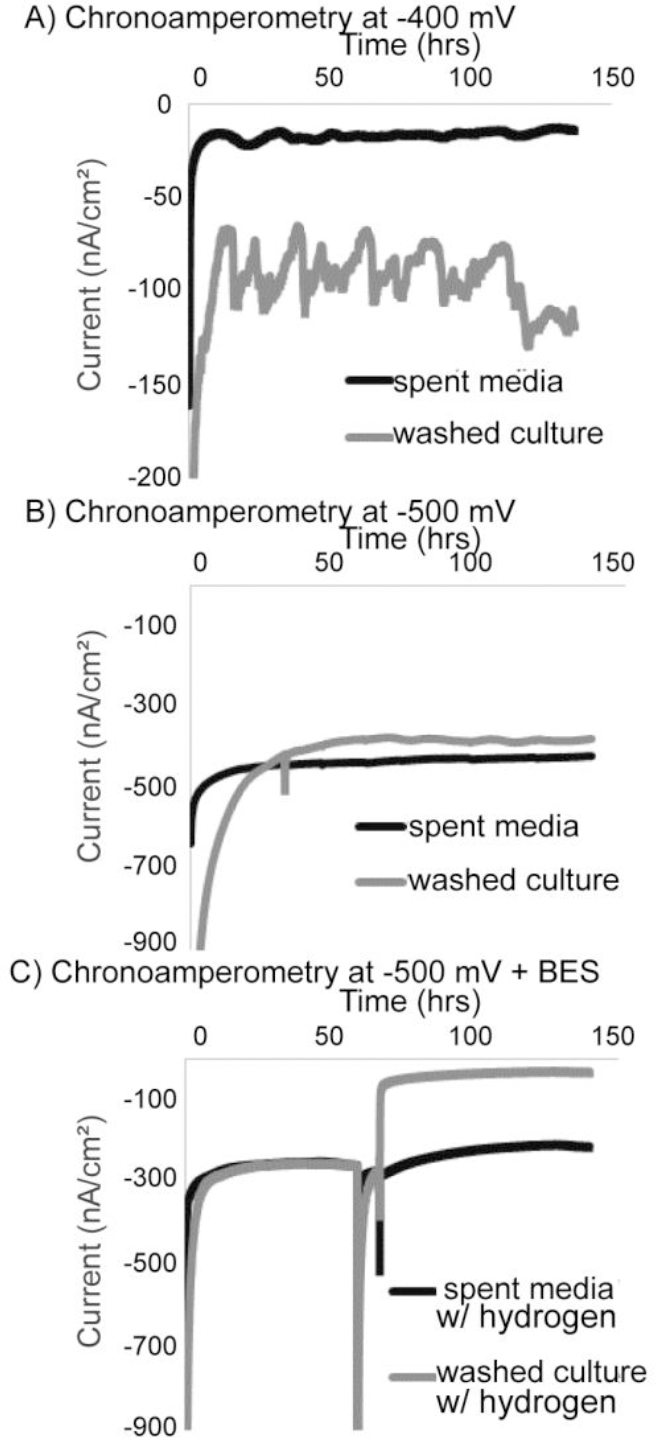
Current consumption occurs in a hydrogenase deletion mutant (Δ*hyd*) across a -400 to -500 mV range and is inhibited by BES. Representative current profiles for the *M. barkeri* mutant (Δ*hyd*) (described in (27)) including experiments with cells (washed culture) and the cell growth media only (spent media) at both -400 mV (A) and -500 mV (B). Changing the hydrogen concentration to include an 80/20 % mix of hydrogen and carbon dioxide did not alter the current consumption, though addition of BES at 18 hours demonstrated inhibition in the reactors containing cells (C).

Cathodic current generation in preparations lacking cells (spent media) of the hydrogenase deletion mutant was observed to be consistently less than that in the washed cells experiments (Figure 3A & B). Additionally, no methane or hydrogen production was observed in the spent media only experiments from the hydrogenase deletion mutant. The lack of hydrogenases, and consequently lack of hydrogen formation in these systems, was further confirmed by the observation that changing the headspace composition of these reactors to a high hydrogen concentration (80/20 H_2_/CO_2_) did not impact cathodic current production in the spent media or washed cell incubations of the hydrogenase mutant (Figure 3C, first 10 hours). To confirm that the electron uptake currents observed in the washed cell experiments were the result of methanogen activity on electrodes, we used 2-bromoethanesulphonate (BES) as an inhibitor of methanogenesis. The addition of 7 mM BES resulted in a dramatic decrease in electron uptake in the washed cell incubation of the hydrogenase deletion mutant, but not in the spent media incubation from the deletion mutant (Figure 3C). This further supports that the observed cathodic electron uptake feature is coupled to methanogenesis.

### Hydrogenase-independent electron uptake feature observed in M. barkeri hydrogenase deletion mutant

Cyclic voltammetry in washed cultures of the *M. barkeri* hydrogenase deletion mutant demonstrated a single low potential electron uptake feature consistent with the feature previously observed for wild type *M. barkeri* cultures at -509 mV vs. SHE (Figure 4A, Figure 2). The deletion mutant experiments were compared with cell free or spent media conditions, where cell cultures were removed via centrifugation. Notably, this feature was not observed in the spent media experiments supporting that this feature is cell-associated rather than extracellular enzyme based (Figure 4A). Cyclic voltammetry confirms that electron uptake in both the spent media (Supplemental Figure S2) and washed cell experiments (Figure 4B) was not altered by addition of 80/20 H_2_/CO_2_, suggesting that hydrogen formation is not occurring in these reactors nor is it affecting current and methane production. This supports the hydrogen and hydrogenase-independent nature of the low potential electron uptake feature observed in *M. barkeri*. The lack of a similar feature in the spent media experiments also supports that the electron uptake feature observed is cell-associated rather than due to a free extracellular enzyme.

**Figure 4.**
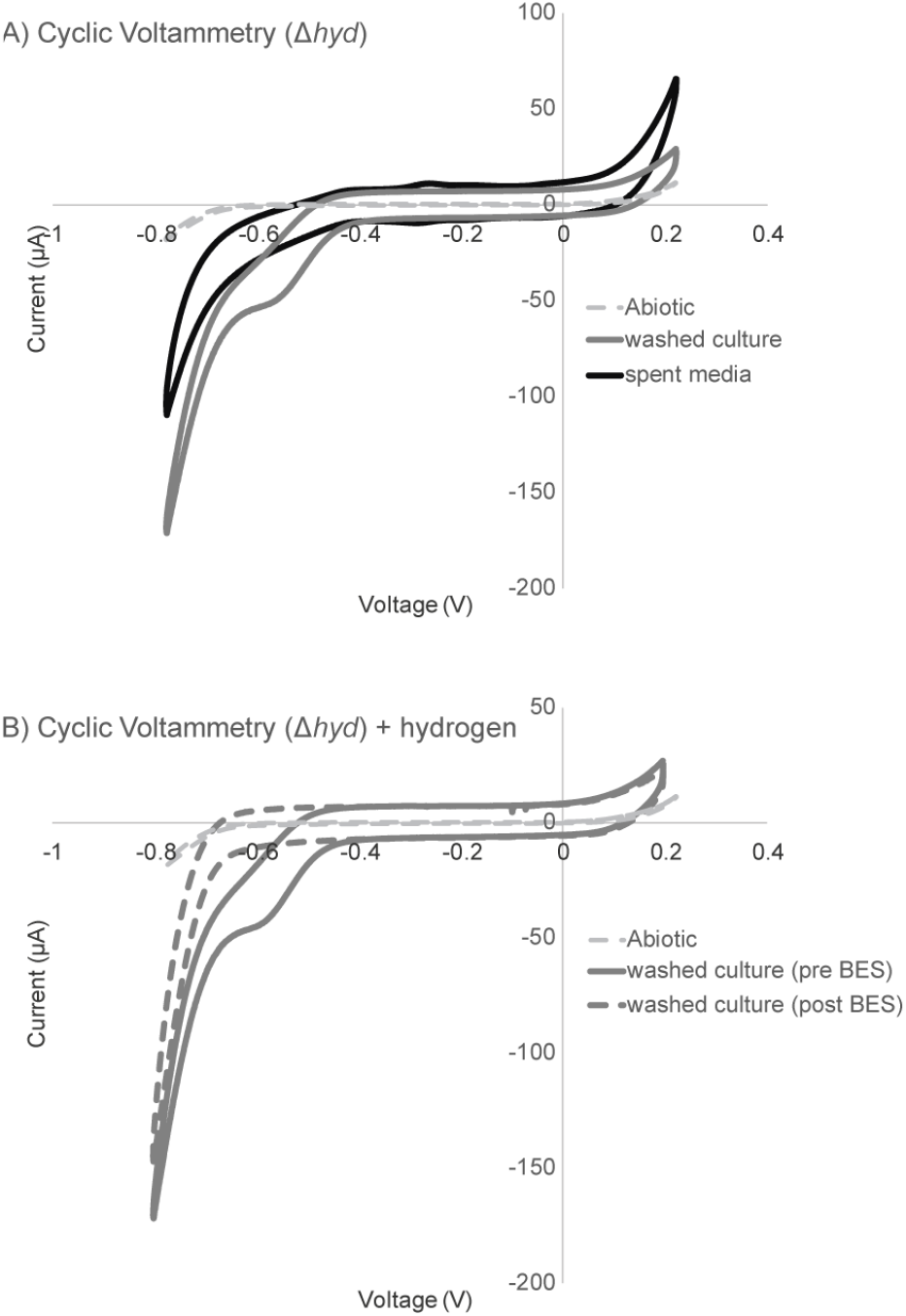
Electron uptake profile of hydrogenase deletion mutant (Δ*hyd*) is similar to *M. barkeri* wild-type washed culture experiments, and is diminished in the presence of BES. Cyclic voltammetry (1 mV/sec scan rate over a -800 to 200 mV range) showing current-voltage relationship for washed cultures of the Δ*hyd* mutant compared with spent media from the Δ*hyd* mutant and blank media abiotic controls (A). Comparing washed culture experiment cyclic voltammetry, hydrogen addition (head space of 80/20 H_2_/CO_2_) results in no change in the electron uptake features observed, however the addition of 7 mM BES abolishes the electron uptake feature (B).

With the addition of BES to washed cells, the formerly observed electron uptake feature is no longer present (Figure 4B). However, BES addition did not alter electron uptake in the cell-free spent media controls (Supplemental Figure S2). This suggests a physiologic linkage between the low potential electron uptake and generation of methane by *M. barkeri.* Inhibition by BES was rapid, occurring in 10-15 minutes (Figure 3C). In other methanogenic electrode systems, a slow onset to BES inhibitions was observed as was a corresponding spike in hydrogen (25). In this alternate case, the inhibition of current was likely linked to product inhibition or specifically the accumulation of hydrogen inhibiting hydrogenase activity. The rate of this inhibition (previously observed to occur at slower time scales of 10+ hours (25)), compared with our observations of a very rapid inhibition helps further distinguish this free extracellular enzyme independent and putatively direct mechanism of microbe-cathode interaction.

### M. barkeri cells are in direct contact with carbon electrodes

To investigate the potential for direct cell-electrode interactions, *M. barkeri* cells from washed culture experiments were stained with NanoOrange and visualized by fluorescence microscopy (Figure 5). Direct contact between cell surfaces and carbon cloth electrode fibers could be observed, with attached cells similar in size and shape (1-2 µm cocci) to previously described planktonic *M. barkeri* cells (26). Similar patterns of cell attachment were also observed using scanning electron microscopy (Supplemental Figure S3). In the spent media controls, intact cells were not observed, but proteinaceous material stained with NanoOrange appeared to attach to the electrodes (Figure 5). Combined with previous observations these data provide evidence for potential direct cell-electrode contacts, providing further support for putatively direct and free extracellular enzyme independent interactions.

**Figure 5.**
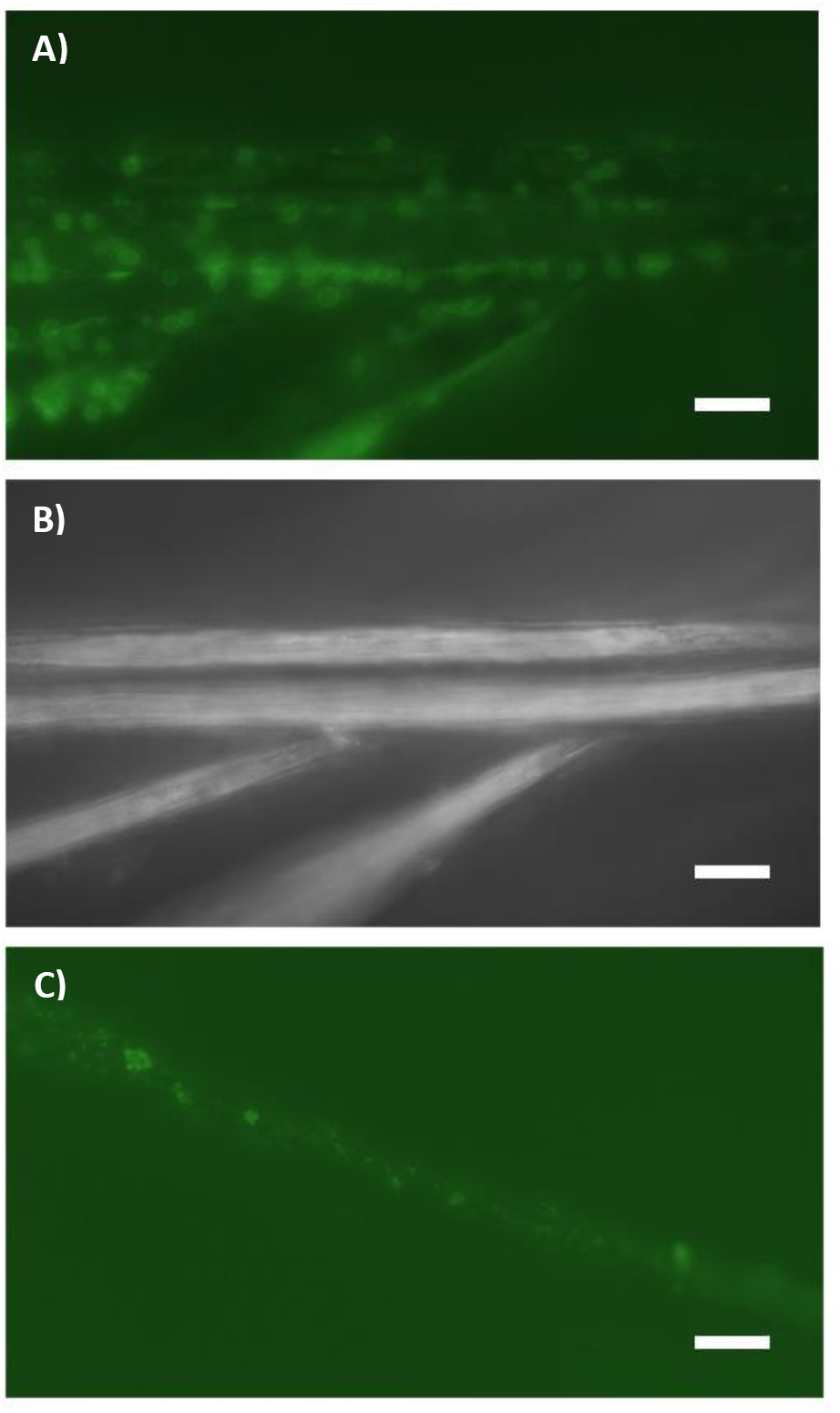
*M. barkeri* attach to carbon cloth fibers of electrode. Experiment treated with NanoOrange^®^ protein stain in washed cell experiments (A) to highlight protein of the cells S-Layer in contact with the carbon fibers of the carbon cloth electrode, which are highlighted in reflective light image (B). NanoOrange^®^ staining of spent media extracts highlights attachment of protenateous material to carbon fibers in spent media only experiments (C).

## Discussion

The data presented here demonstrate the ability of *M. barkeri* to perform electron uptake from cathodes using at least two pathways, that also allow coupling this current to methane production. The higher potential pathway (midpoint potential -301 mV vs SHE) is consistent with the mechanism recently described in *Methanococcus maripaludis*, where attachment of cell-derived free enzymes to carbon electrodes results in the formation of electron donors for methanogenesis (predominantly formate and hydrogen) (18). Here, we provide electrochemical (cyclic voltammetry) data that support this observation of free extracellular enzyme-mediated activity. Specifically, the similarity in redox potential between the electrode oxidation and electrode reduction sweeps suggests a tightly coupled electron transfer event similar to those observed in thin film or protein film voltammetry (19, 20). This mode of electrochemical activity does not require cells to contact an electrode, and seems unlikely to explain the mechanism of direct interspecies electron transfer that has been described previously in microbial consortia (17).

We also detected a previously undescribed hydrogen-independent and free extracellular enzyme-independent mechanism of methane-linked electron uptake on electrodes. The redox potential of this pathway is distinctly lower than those observed for hydrogenases. While this lower potential (electron uptake potential -509 mV vs SHE) presents even more favorable energetics for hydrogen evolution, we directly tested the involvement of hydrogenases by capitalizing on the reversibility of hydrogenases—using elevated hydrogen concentration as a means to inhibit hydrogen formation and drive hydrogen reduction. In addition, we performed experiments using an *M. barkeri* deletion mutant lacking all known methane-linked hydrogenases, that maintained this low potential (-509 mV vs SHE) electron uptake coupled to methanogenesis despite deletion of hydrogenases. Previous work in a *M. maripaludis* hydrogenase deletion mutant demonstrated the ability of extracellular formate dehydrogenases to yield cathodic current through formate production (18). *M. barkeri*, however, has not previously been shown to grow on formate and was not capable of growth using formate under our experimental conditions. Formate addition to our electrochemical experiments resulted in neither stimulation nor inhibition of the electrochemical activity observed, confirming that formate dehydrogenases are unlikely to be playing a role (data not shown).

The difference in biochemistries between *M. maripaludis* and *M. barkeri* likely underlie the differences observed in the predominant mode of electron uptake observed in these electrochemical systems. *M. maripaludis* falls phylogenetically in the clade of methanogens without cytochromes, while *M. barkeri* falls within the clade containing cytochromes. Though several steps in the biochemical pathway between these clades are conserved, a few steps are distinct and result in fundamental differences in the amount of energy captured, as well as the overall reaction rate of methane formation (6). The observed low potential electron uptake feature described in this work may be another example of a biochemical distinction between these two groups. It is feasible that the ability to transfer an electron from a solid phase substrate (electrode) or directly from another cell are due to a related phenomenon as has been observed in these electrochemical systems. In this case, the -509 mV potential at which electron uptake was observed suggests an energetic advantage for cells using direct interspecies electron transfer. Obtaining electrons at a lower redox potential than hydrogen could be energetically beneficial for regenerating reduced ferredoxin required in methanogenesis, mitigating the need for consuming ion motive force. Though more evidence is required to confirm yield, these data suggest a possible ecologic trade-off for methanogens using direct interspecies electron transfer compared with interspecies hydrogen transfer. As of yet, only members of the cytochrome containing methanogen clade have been shown to participate in hydrogen/formate independent syntrophies with organisms such as *G. metallireducens*.

Though this work provides support from the methanogen perspective for direct interspecies electron transfer, it also raises some questions about the potential bioenergetics and ecological trade-off for different strategies of interspecies electron transfer. Future work will continue to investigate this novel mode of electron uptake, its kinetic properties, and its biophysical basis.

## Materials and Methods

### Bacterial Culturing Conditions

The strains of *Methanosarcina barkeri* used have been described previously (24, 27). In brief, a background strain containing the *Δhpt∷ΦC31int-attP* promoter fusion was utilized for wild-type experiments as this strain was utilized for genetic manipulations (28, 29). A hydrogenase deletion mutant with the Ech, Vhu, and Fhr hydrogenases removed was utilized for delta-hydrogenase experiments (24). Each strain was provided by William Metcalf at the University of Illinois at Urbana-Champaign. All strains were grown pre-electrochemical studies in a high salt medium using 5 mM DTT as a reductant (26), at 30°C with 100 mM methanol as the sole methanogenic substrate (i.e. methane via methanol disproportionation). The same high salt media was used for electrochemical experiments, with the exception that the additions of vitamin and mineral mixes, 0.001% resazurin, 5 mM DTT and MeOH were omitted. This media was used in both chambers of the H-cell.

Prior to electrochemical tests, cells were grown to mid log phase on methanol. A ten percent dilution of this biomass was added to each electrochemical chamber for each experiment. For growth culture incubations, 10 mLs of culture were added directly to the working electrode chamber of each reactor. In the case of washed culture incubations (both open circuit and poised potential experiments), approximately 10 mLs of culture were centrifuged at 9000 x g for 10 min in an anaerobic chamber. Cell pellets were resuspended in reactor media and added to respective H cells. Spent media only experiments were performed with 10 mL of media centrifuged at max speed 13000 x g for 20 min as filtering extracts appeared to diminish extracellular protein concentration or activity.

### Electrochemical Reactors

H-cell reactors (Adams and Chittenden Scientific Glass, Berkeley, CA) equipped with two sampling ports in the 150 mL working electrode chamber were utilized for methanogen electrochemical experiments. A Nafion^®^ 117 proton exchange membrane separated the working electrode from counter and reference electrode processes. The membrane was secured via two o-rings and attachment of a vacuum clamp (set up described previously (18, 30)). Ethanol treated carbon cloth attached (∼2 x 3 cm rectangle) to a titanium wire (31) was utilized for a working electrode with the exception of experiments for electron microscopy, where a graphite plate was used for ease of identifying the cell-electrode interfaced. Titanium wire attached to a 4-cm platinum wire was utilized for the counter electrode. Ag/AgCl reference electrodes were purchased from Basi inc. Reference electrodes were calibrated to a standard un-used reference before each experiment and stored in a 3M KCl solution between experiments.

### Electrochemical Conditions

Electrochemical experiments were performed using a CH1010 potentiostat (CHInstrument, TX). Amperometry experiments were run at poised potentials of -300 to -500 mV vs. SHE. Cyclic voltammetry was run over a potential window from +200 to -800 mV using a scan rate of 1 mV/sec unless otherwise specified. First derivatives were calculated using the Origin 61 software package (OriginLab, MA).

### Gas Analysis

Methane production was calculated from head space analysis using a flame ionizing detector (FID) and thermal conductivity detector (TCD) equipped gas chromatograph—a Shimadzu Gas Analyzer (Shimadzu)—as previously described (32). Gas concentrations were calibrated using a 1% and 0.5% standard gas mix (Restek, PA).

### Protein Quantification

Protein content for this work was analyzed using the NanoOrange^®^ protein quantification kit. In short, cell pellets or carbon cloth electrode samples were boiled in 0.5 M NaOH for 10 min. Dilution were quantified to ensure sample analysis within the 10 ng/mL to 10 µg/mL range. Fluorescence was quantified using the Optifluour plate reader.

### Visualization of electrode biomass

Electrode samples were fixed using 2.5% glutaraldehyde in a 25 HEPEs buffer pH 7.5 for 30 min. Post fixation samples received six to ten washes with 50 mM HEPSs buffer at pH 7.5. NanoOrange^®^ protein stain was used to visualize cells using fluorescence microscopy per manufacturers protocol. For scanning electron microscopy (SEM), electrodes were dehydrated in ethanol and then critical point drying using hexamethyldisilazane (HMDS) was used prior to visualization (33).

## Acknowledgements

Annette Rowe was funded in part by a NASA Astrobiology Institute (NAI) postdoctoral fellowship as part as the NAI Life Underground team, as well as by the Innovation Fund Denmark Electrogas project. This work is NAI-LU contribution #. Work in the El-Naggar laboratory was supported by the Division of Chemical Sciences, Geosciences, and Biosciences, Office of Basic Energy Sciences of the US Department of Energy through grant DE-FG02-13ER16415 as well as a Dimensions of Biodiversity grant funded by the NSF (NSF #).

## Supplemental Information for: Methane-linked mechanisms of electron uptake on cathodes in *Methanosarcina barkeri*

**Figure S1.**
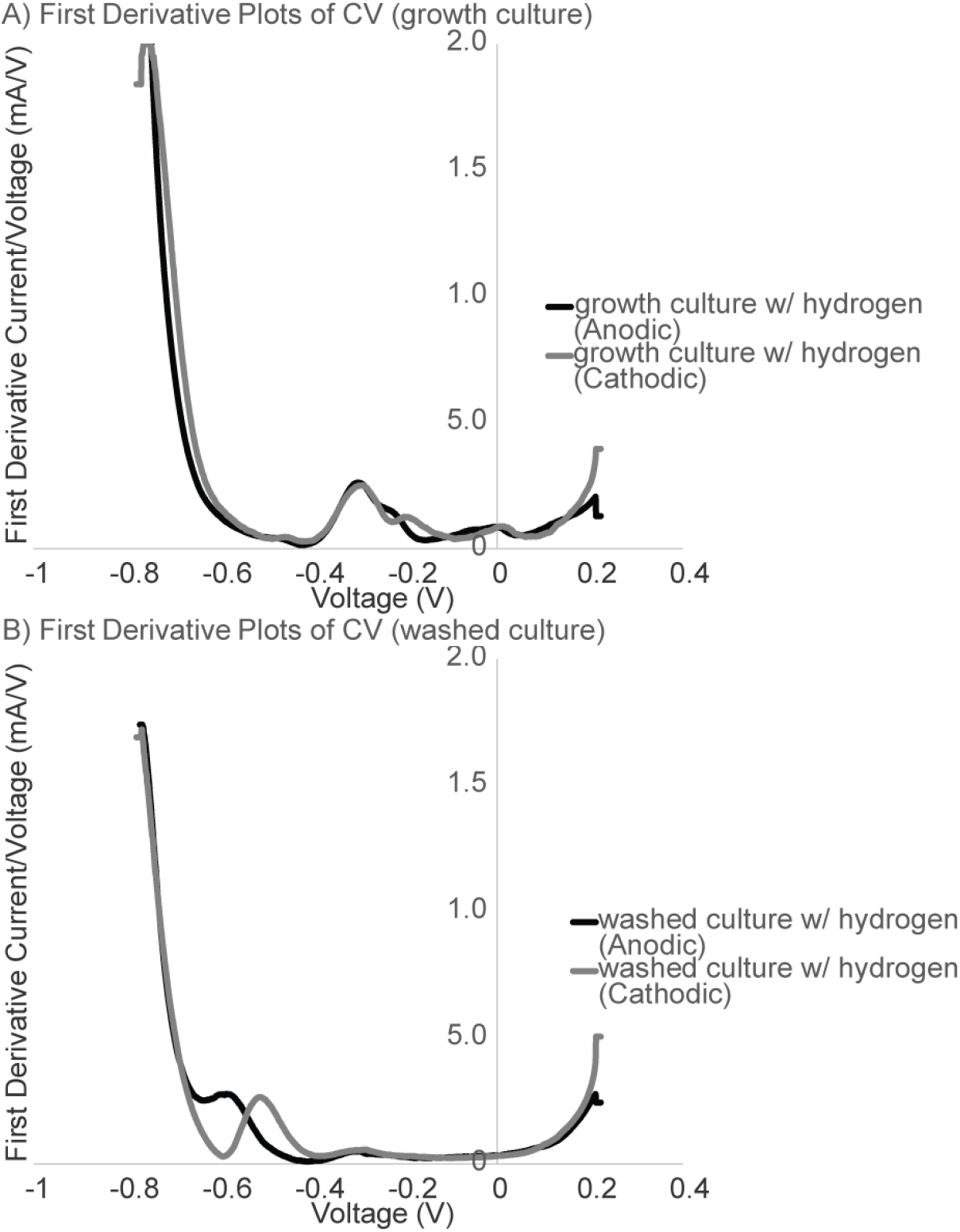
Dominant electron uptake feature in spent media + cells experiments resemble an electrode bound enzyme process. First derivative plots of anodic (negative to positive) and cathodic (positive to negative) sweeps from cyclic voltammetry of growth culture (A) and washed culture (B) experiments. Cyclic voltammetry data shown in Figure 2D of main text.

**Figure S2.**
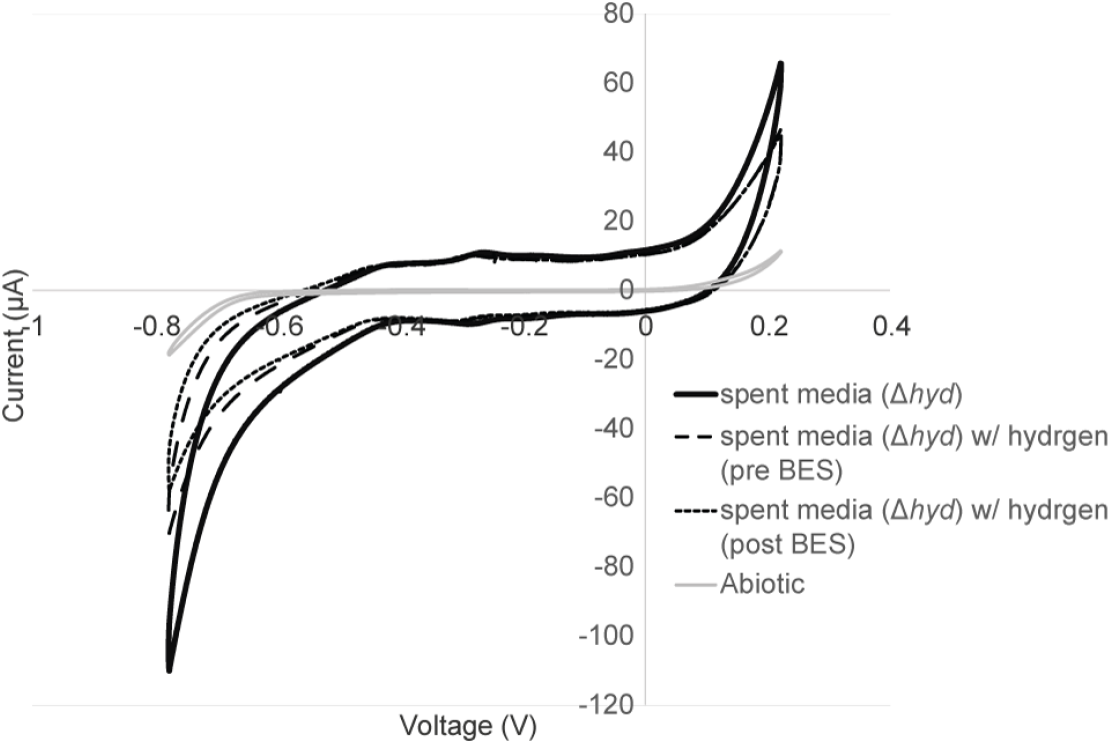
Spent media of hydrogenase deletion mutant shows little to no change with addition of hydrogen and BES. Cyclic voltammetry (1 mV/sec scan rate over a -800 to 200 mV range) showing current-voltage relationship for spent media only controls under experimental conditions with a nitrogen and carbon dioxide atmosphere (80/20 N_2_/CO_2_), a hydrogen and carbon dioxide atmosphere (80/20 H_2_/CO_2_) and with 7 mM BES added.

**Figure S3.**
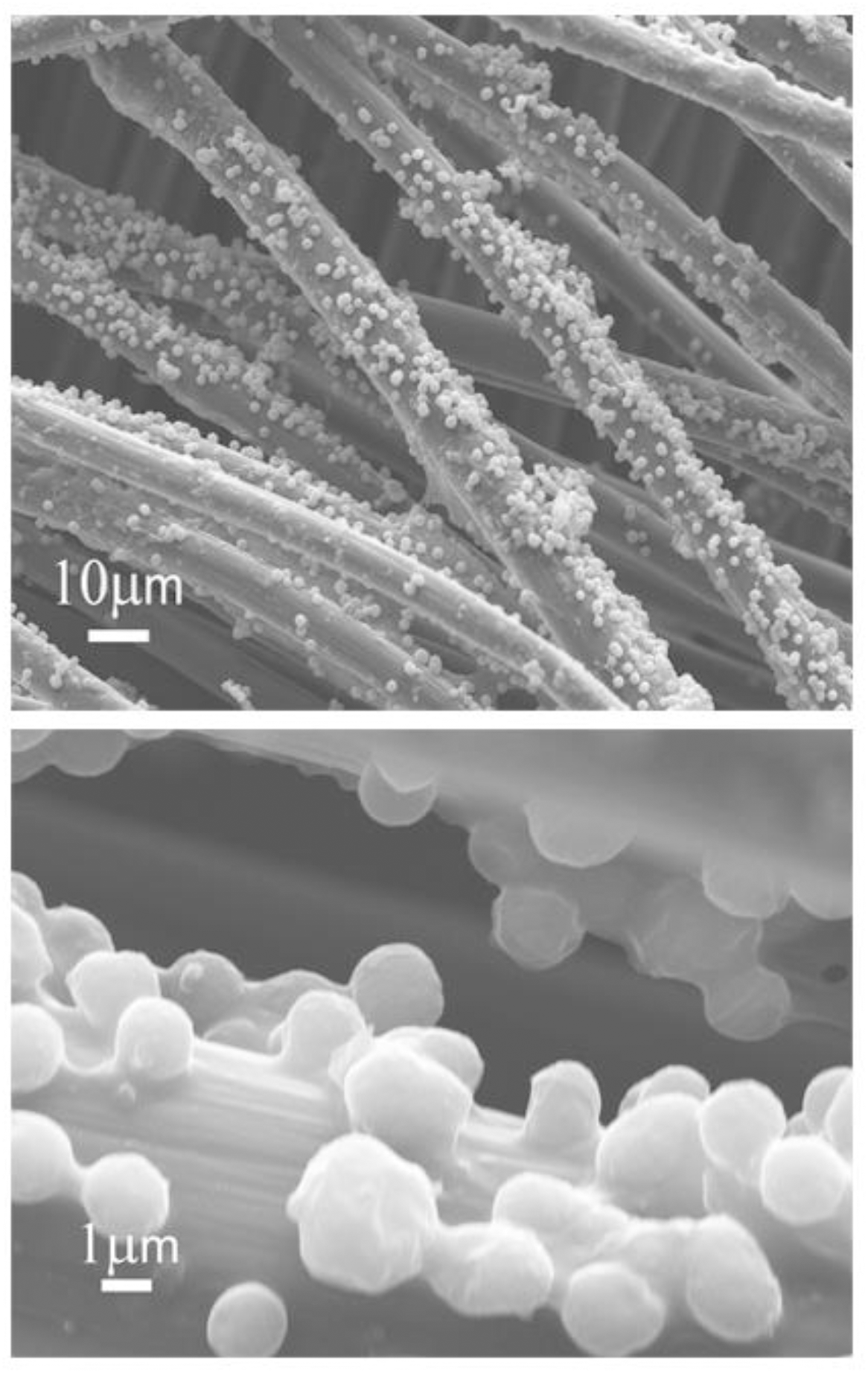
Methanogen cell attachment to carbon fibers of electrode shown via scanning electron microscopy. Images taken of wild-type washed culture experiments poised at -450 mV.

